# Connectomic Assessment of Injury Burden and Longitudinal Structural Network Alterations in Moderate-to-severe Traumatic Brain Injury

**DOI:** 10.1101/2021.04.20.440635

**Authors:** Yusuf Osmanlıoğlu, Drew Parker, Jacob A. Alappatt, James J. Gugger, Ramon R. Diaz-Arrastia, John Whyte, Junghoon J. Kim, Ragini Verma

## Abstract

Traumatic brain injury (TBI) is a major public health problem. Caused by external mechanical forces, a major characteristic of TBI is the shearing of axons across the white matter, which causes structural connectivity disruptions between brain regions. This diffuse injury leads to cognitive deficits, frequently requiring rehabilitation. Heterogeneity is another characteristic of TBI as severity and cognitive sequelae of the disease have a wide variation across patients, posing a big challenge for treatment. Thus, measures assessing network-wide structural connectivity disruptions in TBI are necessary to quantify injury burden of individuals, which would help in achieving personalized treatment, patient monitoring, and rehabilitation planning. Despite TBI being a disconnectivity syndrome, connectomic assessment of structural disconnectivity has been very scarce. In this study, we propose a novel connectomic measure that we call network anomaly score (NAS) to capture the integrity of structural connectivity in TBI patients by leveraging two major characteristics of the disease: diffuseness of axonal injury and heterogeneity of the disease. Over a longitudinal cohort of moderate-to-severe TBI patients, we demonstrate that structural network topology of patients are more heterogeneous and are significantly different than that of healthy controls at 3 months post-injury, where dissimilarity further increases up to 12 months. We also show that NAS captures injury burden as quantified by post-traumatic amnesia and that alterations in the structural brain network is not related to cognitive recovery. Finally we compare NAS to major graph theory measures used in TBI literature and demonstrate the superiority of NAS in characterizing the disease.

## 1. Introduction

Traumatic brain injury (TBI) is a global public health problem with 69 million new cases estimated to occur worldwide each year [14r]. Primarily caused by motor vehicle accidents, falls, and sports concussions, TBI has claimed more than fifty thousand lives in the US alone in 2014 [9], and frequently leads to long-term disabilities [26]. A major characteristic of TBI is the shearing of axons across the white matter, induced by external mechanical forces. Diffuse axonal injury (DAI), as it is called, causes disruptions in the connectivity between brain regions across the network [1,22], leading to cognitive deficits [15] that often require rehabilitation for recovery [11]. Traumatic brain injury is heterogenous in many dimensions including cause, mechanism, and severity of injury, as well as recovery rate and burden of chronic symptoms [39,43]. In treatment and rehabilitation planning, heterogeneity of TBI poses a big challenge that makes subject specific approaches necessary [24,64]. Network level analysis of connectivity disruptions in TBI, therefore, is necessary to provide measures quantifying injury burden of individuals, which would help in achieving personalized treatment, patient monitoring, and informing the patient and caregivers regarding the potential long term progression of the disease [24,72].

Advancements in neuroimaging within the last decades have enabled analysis of connectivity disruptions in TBI with modalities such as functional [23,44] and structural MRI [27,38,73]. Diffusion MRI (dMRI), a structural MRI method measuring the diffusion of water molecules in the tissue, has especially been promising in the analysis of TBI as it has been shown to be sensitive to axonal injury at a microstructural level, that is not captured well in conventional MRI [30,41]. Most of the dMRI based studies investigate axonal injury either locally in isolated brain regions [59] or across certain white matter tracts [68], by using dMRI measures such as fractional anisotropy or cortical thickness [25,30]. Analyses involving such microstructural measures, however, fall short in capturing the impact of TBI on overall network topology.

Analysis of structural connectomes, that is, connectivity maps derived from dMRI data quantifying connections between brain regions, enables evaluation of the brain as a network [63]. Despite TBI being considered as a ‘disconnection syndrome’ due to damaged structural pathways connecting brain regions [22], analysis of structural connectivity disruptions and longitudinal change in network organization is surprisingly scarce [28,34]. The majority of studies investigating structural connectivity in TBI utilize graph theoretical measures, reporting increase in shortest path length [34] and small-worldness [74], and decrease in global efficiency, clustering coefficient [52], betweenness centrality, and eigenvector centrality [15]. While such measures provide insights into the mechanisms of change of the brain’s network structure in TBI, each measure captures a specific aspect of connectivity alteration in the network, which are limited in capturing the overall topological change representing injury burden [6,12,52]. As they are mathematical constructs that are defined for networks at large without any special consideration for brains, interpretation of ensuing results poses further challenges. Additionally, in the absence of a hypothesis that defines the nature of TBI induced change in network topology, it is common to explore a large set of graph theoretical measures that are available in the literature to find those that would demonstrate statistical significance with the data. This exploratory approach, however, suffers from multiple comparison issues [50], affecting TBI studies more than other neuroscientific research due to small sample sizes in the domain. Hypothesis driven studies that suggest markers for TBI by taking the characteristics of the disease into account, on the other hand, are very limited [35,61], and longitudinal analysis of network level change in moderate-to-severe TBI is still lacking.

In this study, we propose a measure that we call network anomaly score (NAS) to capture the integrity of structural connectivity in moderate-to-severe TBI patients by leveraging two major characteristics of the disease, that are, diffuseness of the injury and the heterogeneity of the disease. Diffuseness of the injury can be best captured by a connectome-level measure that is sensitive to the global effects of local connectivity disruptions. Heterogeneity of the disease, on the other hand, can be best captured by a normative measure that compares each patient with a reference healthy control sample. Taking a graph matching based approach, we define NAS as the overall network similarity of moderate-to-severe TBI patients relative to a healthy control sample. We hypothesize that NAS captures the injury burden of individuals with TBI, which we test by calculating correlation between NAS and post-traumatic amnesia scores of patients. We evaluate our measure on a cohort of 34 patients with moderate-to-severe TBI, who underwent dMRI and cognitive assessment at 3, 6 and 12 months post-injury, as well as 35 age- and sex-matched healthy controls. In our analysis, we investigate cross-sectional and longitudinal relationships between the NAS and injury severity, as well as cognitive outcome. We also investigate longitudinal changes in network topology of patients relative to controls as quantified by NAS, and evaluate its relationship with the change in cognitive scores over time. Finally, we compare NAS with standard graph theoretical measures that are commonly reported in TBI literature, in their relationship with injury severity and cognitive outcome.

## 2. Materials and Methods

### 2.1. Participants

The data used in this study was acquired as part of a larger project investigating the neuroimaging correlates of functional recovery after diffuse TBI (PI: JJK). All participants provided informed consent directly or via a legally authorized representative. Study procedures were approved and overseen by the Institutional Review Board at the Moss Rehabilitation Research Institute, Elkins Park, Pennsylvania, and the University of Pennsylvania. The cohort investigated in this study consists of 40 participants with moderate-to-severe TBI and 35 healthy controls (HC) [61]. Inclusion criteria for TBI participants were being in the age range 18 to 64 and diagnosis of non-penetrating moderate-to-severe TBI, indicated by at least one of the following: *i*. Glasgow Coma Scale score less than 13 in the emergency department (ED not due to sedation, paralysis, or intoxication), *ii*. documented loss of consciousness for more than 12 hours, *iii*. prospectively documented PTA greater than 24 hours. Exclusion criteria for TBI participants were *i*. history of prior TBI, CNS disease, seizure disorder, schizophrenia, or bipolar disorder, *ii*. history of long-term abuse of alcohol or psychostimulants that could have resulted in neurologic sequelae, *iii*. pregnancy, *iv*. inability to complete MRI scanning due to ferromagnetic implants, claustrophobia, or restlessness, *v*. nonfluency in English; or *vi*. a level of disability preventing completion of testing and scanning by 3 months post-injury. TBI participants with total estimated volume of focal intraparenchymal lesions larger than 5 cm^3^ for subcortical lesions and larger than 50 cm^3^ for cortical lesions were also excluded to ensure that the TBI was predominantly diffuse. Healthy controls recruited were comparable in age, sex, and education to TBI subjects. Exclusion criteria for HCs were the same with TBI participants with the addition of exclusion for any history of TBI resulting in alteration or loss of consciousness.

Cognitive assessment and dMRI scans were obtained for HCs once and for patients three times at approximately 3, 6 and 12 months post-injury. Imaging data was not available for some of the patients at certain time points due to either the patient not attending a follow up session or the data being removed from the dataset because of MRI quality issues such as segmentation problems arising from lesion in the brain. In our analysis, we removed 6 patients from the dataset whose imaging data failed the imaging quality assessment (QA) at 3 months post-injury, leaving 34 patients (12 f) to be analyzed for the study. Among these patients, 27 (10 f) had dMRI data available at 6 and 12 months. We note that dMRI data of only 22 (8 f) of the patients had passed the imaging QA at all three time points. In order to increase the power of the analysis, we used all patient data available at follow up sessions rather than doing the analysis with the patients that have data at all time points. Demographics of the participants are detailed in Table 1.

**Table 1.**
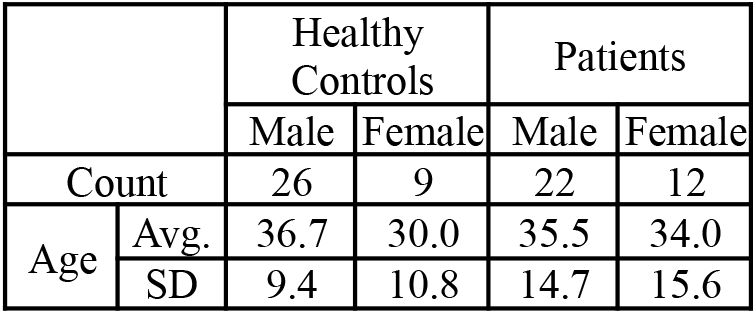
Demographics of the moderate-to-severe TBI dataset with healthy controls.

### 2.2. Data acquisition, preprocessing, and connectome construction

Structural MRI scans were acquired on a Siemens 3T TrioTim whole-body scanner with an 8-channel array head coil (single-shot, spin-echo sequence, TR/TE = 6500/84 ms, b=1000 s/mm2, 30 directions, flip angle = 90°, resolution = 2.2×2.2×2.2 mm). High-resolution T1-weighted anatomic images were also obtained using a 3D MPRAGE imaging sequence with TR = 1620 ms, TI = 950 ms, TE = 3 ms, flip angle = 15°, 160 contiguous slices of 1 mm thickness, FOV = 192×256 mm^2^, 1NEX, resolution = 1×1×1 mm. T1 images were preprocessed using the FreeSurfer 5.3.0 recon-all pipeline (http://surfer.nmr.mgh.harvard.edu) [18] and registered to the FA using rigid followed by deformable SyN registration in ANTs [2] with the deformation constrained to the anterior-posterior direction to correct for the EPI distortions in the dMRI. 86 regions of interests from Desikan atlas [13] were extracted to represent the nodes of the structural network. Five-tissue-type images for anatomically constrained tractography (ACT) [60] were created from Freesurfer outputs. 500 seeds for tractography were placed at random inside each voxel of the mask of the grey-matter white-matter interface (GMWMI). Probabilistic tractography was performed in mrtrix3 [66] using the iFOD2 algorithm [65] with angle curvature threshold of 60°, step size of 1 mm, and minimum and maximum length thresholds of 25 mm and 250 mm, respectively. Connectomes were then generated as an 86×86 adjacency matrix of weighted connectivity values, where each element represents the number of streamlines between regions. Each connectome was subsequently normalized by the GMWMI volume of the individual.

### 2.3. Behavioral and Cognitive Measures

TBI patients underwent behavioral assessment at each time point to yield one behavioral and three cognitive measures. Duration of post-traumatic amnesia (PTA), calculated as the number of days between the TBI and the first of two occasions within 72 hr that the patient was fully oriented, was used as a sensitive behavioral index of the injury severity [4,51]. Full orientation was defined as a score above 25 on the Orientation Log [31], or documentation of consistent orientation for 72 hr in the medical record.

Three cognitive measures were assessed: information processing speed (PS), verbal learning (VL), and executive functioning (EF). We used Processing Speed Index from the Wechsler Adult Intelligence Scale-IV [70] to assess PS and The Rey Auditory-Verbal Learning Test [54] to evaluate VL. A composite score was used for assessing EF to reduce type I error and increase signal-to-noise ratio, which is calculated as a combination of the scores obtained from the following five tests: Controlled Oral Word Association Test [5], Trail Making Test-Part B [53], Color-Word Interference Test, and Digits Backward and Letter-Number Sequencing subtests from the Wechsler Memory Scale IV [71]. We identified the rank of a participant on each individual measure and averaged the ranks across five measures to form the composite executive measure.

### 2.4. Network Anomaly Score

In order to evaluate change of brain’s network organization in TBI patients over time, we consider graph matching [19] to quantify connectomic similarity as it accounts for changes in the overall topology of the network rather than focusing on local changes in individual connections. Previously, we have successfully applied graph matching in deriving similarity between connectomes for quantifying injury severity in TBI patients [46], evaluating subject-wise structure-function correspondence [47], and investigating connectomic stability within and across subjects [48]. In this study, we extend our previous approach by adopting a different use of graph matching to provide a normative connectomic similarity measure.

#### A graph matching based measure to quantify connectomic similarity

Here, we first provide a brief overview of graph matching. Given two graphs *A* and *B* that are deemed to have a similar topology, the aim of graph matching is to find the optimal mapping between the two graphs by assigning each node of *A* to a node of *B* that structurally resembles it the most. Given a cost function *c : A* → *B* → ℝ determining the cost of assigning each node in *A* to a corresponding node in *B*, graph matching can be formulated as a combinatorial optimization problem where the aim is to calculate a one-to-one mapping *f* : *A* → *B* between the nodes of *A* and *B* by minimizing the objective function 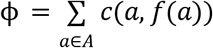 On connectomes, we regarded the cost function *c* as the Euclidean distance between the *k*-dimensional feature vectors of nodes encoding their connectivity signature relative to other nodes in a parcellation with *k* ROIs. We obtained the desired mapping by solving the optimization problem using the Hungarian algorithm [36]. Since brain structure has commonalities across people and the parcellation that yielded graph representations of brains are the same across subjects, we expect the resulting mapping to match nodes of *A* with their corresponding nodes in *B* (i.e., the matching nodes should correspond to the same ROI), which we call a *correct match*. On the other hand, if the connectivity patterns of the nodes vary too much between the two graphs, it would lead to *incorrect matching* of some of the nodes where nodes in *A* will be assigned to nodes in *B* that are not their counterparts. Consequently, we regarded *network similarity (NS)* as the percentage of correct matches relative to total number of nodes (Fig.1.a), with larger values indicating higher similarity. Using this graph matching based measure in quantifying network similarity allows capturing the similarity of overall network organization since matching between the nodes are obtained through the solution to an optimization problem.

**Figure 1.**
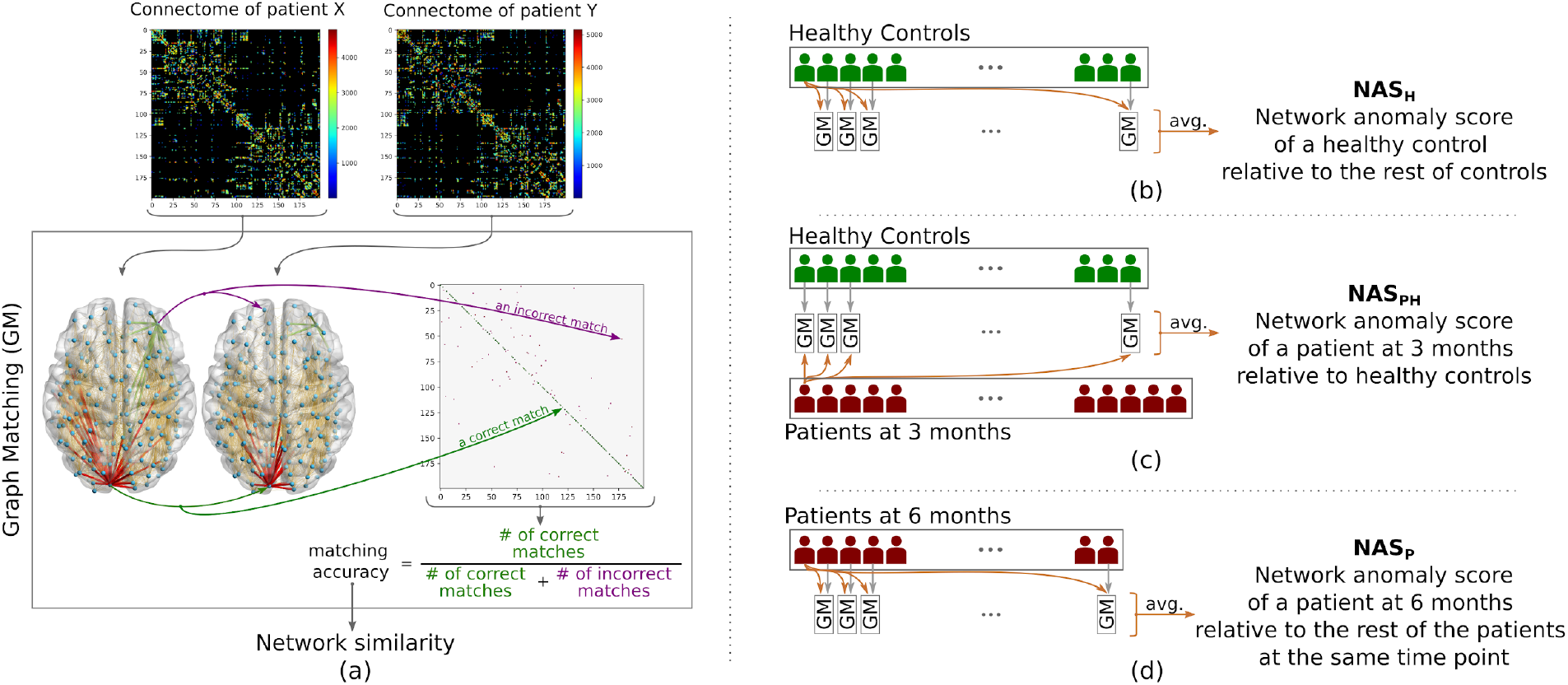
Network anomaly score (NAS) quantifying similarity of structural network organization of a subject’s brain relative to a sample. (a) Taking two connectomes representing the structural connectivity of two subjects as input, the similarity between their graph representation is calculated using graph matching, yielding a binary matching matrix. Similarity between the connectomes are determined as the proportion of nodes which were correctly matched. Using graph matching (GM) as the measure of network similarity, we calculate network anomaly score of (b) each healthy control relative to the rest of the healthy controls (NAS_H_), (c) each patient at a certain time point relative to healthy control sample (NAS_PH_), and (d) each patient relative to the rest of the patients at the same time point (NAS_P_).

#### Normative connectomic similarity: similarity of a subject relative to a sample

Having defined NS as the similarity measure between two connectomes, we next define the network anomaly score (NAS) as a normative measure consisting of the mean NS of the subject relative to the reference sample (Fig. 1.b-d). Taking healthy controls as the reference, we first calculated network anomaly score among them to provide a basis for evaluation (Fig.1.b). We then calculated similarity of patients at a certain time point (such as 3 months) relative to the healthy (Fig.1.c), to quantify trauma induced network alterations in TBI patients. In order to evaluate heterogeneity and the course of relative changes in network topology among patients, we calculated a third anomaly score quantifying similarity of patients relative to the rest of the patients within the same time point (Fig.1.d). For the sake of clarity, we refer to these three scores as NAS_H_, NAS_PH_, and NAS_P_ in the rest of the paper, where subscripts denote NAS among the healthy, NAS of patients relative to healthy, and NAS among the patients, respectively.

### 2.5. Statistical Analysis

#### Group level analysis

In order to evaluate group differences in the structural network organization cross-sectionally, we ran the Mann-Whitney U test between NAS_PH_ (or NAS_P_) and NAS_H_, and the Wilcoxon signed-rank test between NAS_PH_ and NAS_P_. We quantified the amount of change in scores using the following effect size formula:

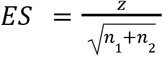

where n_1_ n_2_ are the sample size of groups, and 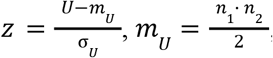 and 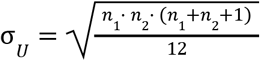, and *U* is the test statistic for the Mann-Whitney U test, whereas 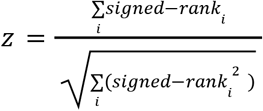 for the Wilcoxon signed-rank test. Effect size is regarded as small if |ES| ≥ 0.1, medium if |ES| ≥ 0.3, and large if |ES| ≥ 0.5.

#### Cross-sectional linear model analysis

In evaluating the relationship between NAS and injury severity or cognitive scores cross-sectionally, we utilized a linear model (LM) format that controls for age and sex as follows:

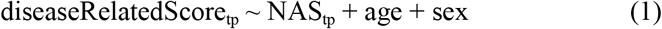

where diseaseRelatedScore is replaced by one of the cognitive scores or PTA, while tp indicates one of 3, 6 or 12 months time points. Analyses were done in R using the nlme package [49].

#### Longitudinal linear mixed effect model analysis

In order to investigate whether the network organization of patients demonstrates a linear change over time when considered altogether, we evaluated the longitudinal change in their network anomaly scores (NAS_PH_ and NAS_P_ are evaluated separately). Since imaging data was not available at all time points for some subjects, we used linear mixed effects (LMEM) analysis with the following model:

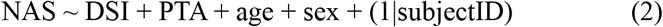

where we estimated NAS as a linear function of the fixed variables days since injury (DSI), PTA, age, and sex, along with the random intercept. Analyses were done in R using the lme4 [3] and lmerTest [37] packages. In our LM and LMEM analysis, we scaled the values of variables. Thus the estimated values of independent variables (e.g. DSI, PTA, age, etc. in eqn. 2) can be interpreted as their correlations with the dependent variable (e.g., NAS in eqn. 2).

#### Analysis of the trajectory of change

In order to evaluate whether there exists a relationship between the cognition and NAS in how they change over time, we calculated the rate of change as the slope of the line connecting measurements between two time points for each score type. We then calculated Pearson’s correlation between resulting terms to report relationships.

### 2.6. Standard graph theoretical measures

In order to further highlight the efficacy of the proposed score in characterizing TBI, we evaluated our moderate-to-severe TBI cohort using standard graph theoretical measures that are cited in the TBI literature. We considered node betweenness centrality, eigenvector centrality, clustering coefficient, small worldness, characteristic path length, global efficiency, and modularity. We used the Python implementation (bctpy, version 0.5.2, https://pypi.org/project/bctpy/) of Brain Connectivity Toolbox [55] to calculate the measures over the connectomes. The statistical analysis for NAS was repeated for each of these graph theory measures individually (see SI.4 for further details on graph theory measures and their analysis).

## 3. Results

### 3.1. Group level analysis of network similarity between patients and controls

In order to evaluate whether the proposed measure captures structural connectivity alterations, we performed a group-level analysis between patients and healthy controls (Fig.2). We observed significantly lower network similarity scores for patients (NAS_PH_) compared to healthy controls (NAS_H_) at 3 months (ES=0.42, p<10^−3^), 6 months (ES=0.41, p=10^−3^), and 12 months (ES=0.59, p<10^−4^). This result shows that the NAS captures TBI induced alterations of the network topology to distinguish structural connectivity of patients from that of healthy controls up to 12 months post-injury.

**Figure 2.**
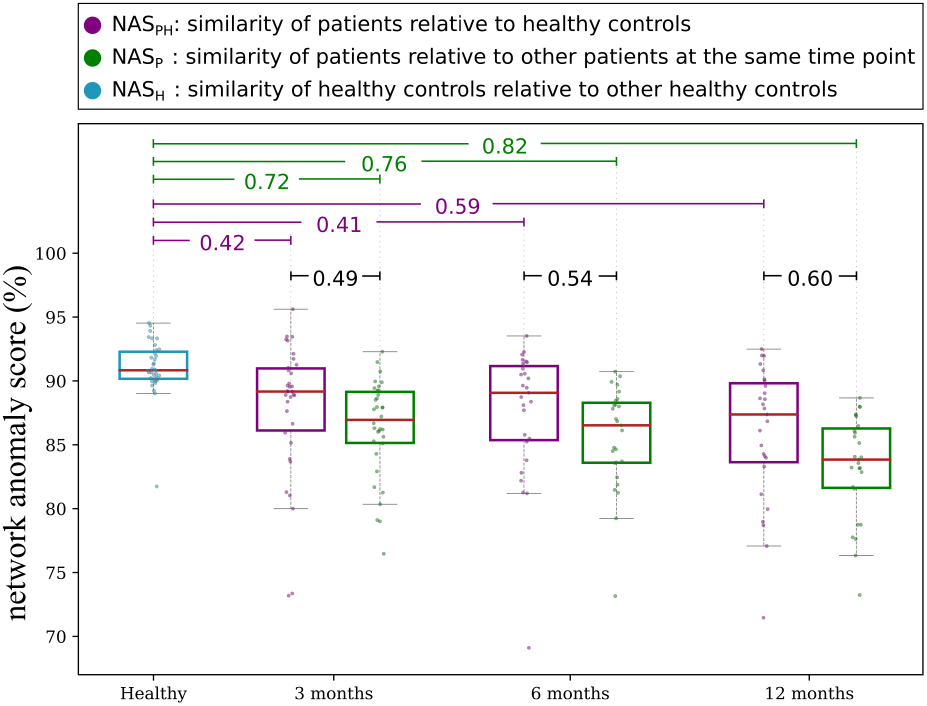
Group level analysis of network anomaly score across patients and controls. We evaluated network anomaly scores (NAS) of patients relative to healthy controls (NAS_PH_), NAS among patients within the same time point (NAS_P_), and NAS among healthy controls (NAS_H_). We observed that network topology of patients is significantly dissimilar to that of the healthy (NAS_PH_ < NAS_H_, purple lines at the top), showing that trauma induced injury introduced alterations across the network. We also observed that NAS_H_ > NAS_P_ with statistical significance (green lines at the top), highlighting a larger variance of network topologies among patients than controls. We then observed that NAS_PH_ > NAS_P_ (black lines at the top), indicating that patients resemble the healthy more than they resemble other patients. These results further show that the heterogeneity of the disease is captured at the structural brain network topology of patients. (Note that lines at the top between pairs of sample groups show effect size for significant group differences with p<0.05, results are FDR corrected)

We then investigated whether NAS captures the heterogeneity of the disease at the network level. We observed that patients had significantly lower within-group network similarity scores (NAS_P_) compared to healthy controls (NAS_H_) at 3 months (ES=0.72, p<10^−6^), 6 months (ES=0.76, p<10^−6^), and 12 months (ES=0.82, p<10^−6^). This result underlines a higher heterogeneity in structural network topology among patients than that among healthy controls, indicating that the injury affecting each patient differently leads to a unique network organization. We also observed a significant group difference between NAS_PH_ and NAS_P_ at 3 months (ES=0.49, p<10^−4^), 6 months (ES=0.54, p<10^−4^), and 12 months (ES=0.60, p<10^−4^), which indicate that network structures of patients resemble that of healthy controls more than they resemble that of other patients.

### 3.2. Relationship between network similarity and injury severity

A significant negative association between PTA and NAS_PH_ was observed (see eqn. 1 for the LM) at 3 (p=0.016, est_NAS_=-0.51), 6 (p=0.016, est_NAS_=-0.48) and 12 months (p=0.016, est_NAS_=-0.52), while no significant association was observed for age and sex (see Table 2). This result indicates that more severely injured patients have lower network similarity in reference to healthy controls.

**Table 2.** Results of fitting a linear model to evaluate the relationship between injury severity (PTA) and NAS_PH_, age, and sex. We note that since the scores were scaled for the LM analysis, the estimated values provided in the table (columns labelled with “est”) indicate correlation of corresponding variables with PTA (see eqn. 1 for LM, p-values are FDR corrected for each variable across three models).

| Time Point | Adj.$R^2$ | $p_{NAS}$ | $est_{NAS}$ | $p_{age}$ | $est_{age}$ | $p_{sex}$ | $est_{sex}$ |
| --- | --- | --- | --- | --- | --- | --- | --- |
| **3 Months** | 0.202 | **0.016** | **-0.509** | 0.810 | -0.083 | 0.270 | 0.568 |
| **6 Months** | 0.223 | **0.016** | **-0.480** | 0.810 | 0.045 | 0.454 | 0.336 |
| **12 Months** | 0.176 | **0.016** | **-0.521** | 0.810 | -0.178 | 0.454 | 0.276 |

### 3.3. Change in network anomaly score over time

Group level analysis of network similarity scores shown in Fig.2 demonstrated an increase in effect size between patients and controls from 3 to 12 months, suggesting that the structural connectivity in patients becomes more unlike healthy controls over time. A further longitudinal analysis of NAS_PH_ using LMEM (eqn. 2) showed that the similarity score is a function of days since injury (DSI), PTA, and age (p_DSI_<10^−3^, p_PTA_=0.004, p_age_=0.019, p_sex_=0.581, Adj. R^2^=0.316), with a negative association (est_DSI_=-0.144, est_PTA_=-0.429, est_age_=-0.33) (see Table SI.1.a for further details). This result indicates that patients become significantly unlike healthy controls in their structural network connectivity as time progresses post-injury up to 12 months.

Repeating the same analysis for network similarity among patients, we observed a significant decrease of NAS_P_ over time (p_DSI_<10^−4^, p_PTA_=0.007, p_age_=0.066, p_sex_=0.662, Adj. R^2^=0.298) with a slope of −0.306 for DSI, indicating a steeper decline when compared to change in NAS_PH_ with a slope of −0.144 (see Table SI.1.b for further details). This result indicates that although patients deviate from “normalcy” as defined by the network topology of the healthy, they do not converge to an alternate normal that would be common among patients either.

### 3.4. Relationship between the network similarity score and cognitive scores

We next investigated whether NAS_PH_ captures information regarding cognitive function. Before evaluating the relationship between NAS_PH_ and cognition, we first did a group level and LMEM analysis of cognitive scores to evaluate their change over time and their relationship with PTA. We observed that the patients perform significantly lower than controls at 3 months for each cognitive score type (Fig. SI.2. top) and that their performance in each category improves over time significantly to reach the level of healthy controls at 12 months (Fig. SI.2. bottom). We also observed a significant negative correlation between each cognitive score and PTA, with verbal learning (VL) having a marginal p-value (see Supplementary SI.2 for further details). These results show the presence of cognitive recovery in patients up to 12 months post-injury and demonstrate that cognitive performance is related to injury severity.

Observing a disparity between cognitive recovery and increasingly abnormal network topology in patients, we evaluated whether there exists a meaningful relationship between the two virtually diametrical trends (Fig.5). Calculating Pearson’s correlation between rates of change of NAS_PH_ and each cognitive score separately, we observed no significant relationship at any of the time intervals (i.e., 3-6 months, 3-12 months, or 6-12 months, p>0.05 for all tests after FDR correction), indicating the lack of an association between cognitive recovery and change in structural connectivity organization of patients.

Despite the lack of a significant relationship between the rate of change in cognitive scores and NAS_PH_, we evaluated whether there exists a relationship between the actual scores. Using an LMEM (see eqn. SI. 3), we observed that executive function (EF) and processing speed (PS) are significantly and positively related with NAS_PH_ and DSI (EF: p_NAS_<10^−3^, p_DSI_<10^−4^, R^2^_m_=0.206, PS: p_NAS_=0.006, p_DSI_<10^−4^, R^2^_m_=0.226) while verbal learning did not reveal any significant relationship with NAS_PH_ (p_NAS_=0.086, p_DSI_=10^−4^, R^2^_m_=0.141) (see Table SI.3 for further details). The positive correlation between NAS_PH_ and EF and PS indicates that patients with structural connectivity more similar to healthy controls demonstrated better cognitive function.

### 3.5. Evaluation of the cohort with standard graph theory measures

In our analysis of graph theory measures, we first evaluated the association between NAS_PH_ and graph theory measures longitudinally and cross-sectionally, and observed no significant relationship (see Tables SI.4.a and SI.4.b, p-values are FDR corrected for multiple comparison correction, see SI.4 for a detailed explanation of the analysis). We then evaluated the association between graph theory measures and PTA using LM (see eqn SI.4.b) showed statistical significance only for node betweenness centrality at 6 months (p_NBC_=0.002, p_age_=0.962, p_sex_=0.984, Adj. R^2^=0.504) (see Table SI.4.c). Finally, we evaluated the association between cognitive scores and graph theory measures using a LMEM analysis (eqn. SI.4.c). After FDR correction, no significant association was observed (see Table SI.4.d).

## 4. Discussion

Traumatic brain injury is considered a disconnectivity syndrome [22] due to the diffuse injury of axons across the brain tissue, leading to structural connectivity disruptions among brain regions. While local microstructural changes in the brain [27,38,73] as well as functional connectivity alterations [23,44] are well studied in TBI, literature focusing on the structural connectivity changes in the brain has been very limited [28]. This small body of work has two main limitations: First, most of these studies utilize standard graph theoretical measures in their analysis, which are limited in capturing the diffuse characteristics of the injury. Second, although cross-sectional studies are abundant, longitudinal analysis of structural changes in the brain’s network topology and its relationship with cognitive function of TBI patients are scarce. In this study, we proposed a novel measure called Network Anomaly Score (NAS) that is tailored to capture the two established characteristics of TBI, the diffuseness of the injury [1] and the heterogeneity of the disease [43]. In a moderate-to-severe TBI cohort, we demonstrated that the NAS captures injury-induced structural connectivity alterations by quantifying the connectivity differences at the network level. This highlighted a significantly different network topology among patients relative to healthy controls. Our results also show that the heterogeneity of the disease is observable in the network topology of the patients as quantified by the NAS. We further observed that the network structure of the patients becomes more unlike that of healthy controls over time, despite cognitive recovery over the same interval. As we did not observe any significant relationship between the change in cognitive scores and the change in network similarity of patients over time, these results highlight a mismatch between structural change and cognitive recovery. Finally, we demonstrated that the NAS captures characteristics of TBI that are not captured by standard graph theory measures as there was no significant association between the NAS and any of the graph theory measures. We also observed that only node betweenness centrality demonstrated a significant association with injury burden at 6 months, and none of the measures showed a significant association with cognitive scores, as the results didn’t survive multiple comparisons correction. Overall, our results point to a new direction of research in the analysis of structural network alterations in TBI, involving similarity measures that are designed to capture the characteristics of the disease such as heterogeneity and diffuse injury.

### 4.1. Overall network similarity of TBI patients relative to the healthy, captures injury induced alterations in the structural connectivity

The negative correlations between PTA and NAS indicate (Fig. 3) that patients with more severe brain injuries (high PTA score) have network topologies that are less like healthy controls (low network similarity score). When considered with the group level differences of network topologies between patients and controls (Fig. 2), these results highlight the efficacy of the NAS in capturing trauma induced alterations.

**Figure 3.**
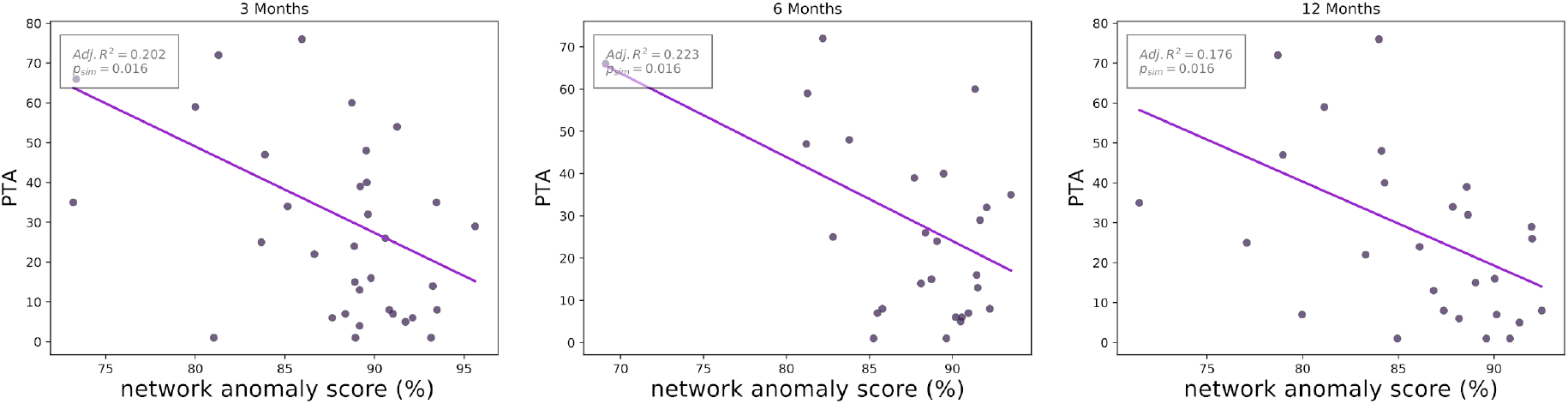
Relationship between network anomaly score and injury severity. Evaluating whether injury severity (PTA) can be described cross sectionally as a function of NAS_PH_, age, and sex using an LM, and we observed a significant relationship between NAS_PH_ and PTA at 3, 6, and 12 months post-injury (p-values are FDR corrected). This result indicates that trauma induced alterations at network topology captures injury severity.

The direct relationship between diffuse axonal injury and the disruptions in structural connectivity among brain regions underlines the potential of a network topological analysis in quantifying injury burden of TBI patients. Interestingly, very few studies in the already limited structural connectivity literature of TBI have evaluated this relationship [6,52]. We were able to identify three studies that considered graph theoretical measures to evaluate network abnormalities of patients and evaluated their relationship with injury severity, two of which reported a lack of a significant relationship in moderate-to-severe adult [6] and pediatric [12] TBI patients, while the third reported a positive correlation for node strength and global efficiency scores [52]. A recent study by our group proposed the Disruption Index of the Structural Connectome (DISC) as a specialized network level score for capturing injury burden on TBI, that demonstrated a significant correlation with injury severity of patients [61]. However, this study was limited in being cross-sectional and the connectivity disruptions being quantified on the basis of edges, rather than at network level.

The lack of significant associations of graph theory measures with PTA and cognitive scores (except for node betweenness centrality at 6 months with PTA) along with lack of a significant relationship between the NAS and any of those graph theory measures, indicate the novelty and superiority of our measure over standard graph theory measures in characterizing TBI. We note that standard graph theory measures are mathematical constructs that are designed to evaluate any graph structure such as social networks or airline route maps, without any specific consideration for brain networks. In the absence of a hypothesis on which measure to use as a biomarker, exploratory analysis that investigates several graph theory measures becomes inevitable. This, however, reduces statistical power of the study due to multiple comparisons correction, which is already limited in TBI studies due to small sample sizes. Interpretation of ensuing results is a further challenge due to measures not being disease specific. Designed specifically to capture well known characteristics of TBI, on the other hand, our proposed measure has two major strengths over standard graph theory measures. First, it focuses on leveraging the diffuse characteristic of the injury by taking a graph matching approach. Since graph matching quantifies similarity through solving an optimization problem, it considers connectivity differences across the network altogether, rather than summarizing connectivity differences on the basis of individual edges. Second, it is a normative score that is calculated relative to healthy controls that leverage the heterogeneity of the disease.

### 4.2. Heterogeneity of the disease is observable in the structural connectivity among brain regions

A major characteristic of TBI is its heterogeneity in various aspects including the cause of initial injury (eg., fall or motor accident), mechanism (eg., direct impact or acceleration/deceleration), pathology (eg., focal and/or diffuse axonal injury), severity (eg., mild, moderate, or severe), ensuing cognitive deficits, and treatment of the disease [29,39] as well as outcomes in cognitive recovery [43]. Lower NAS of patients relative to controls show that network topology of TBI patients differs from healthy control population at varying degrees (Fig. 2). Network similarity among patients being even lower than their similarity relative to healthy controls further supports the previous result, highlighting that injury affects each patient in different ways, potentially due to heterogeneity of the disease in its etiology, mechanism, and severity. In combination, these results demonstrate for the first time in the literature that the heterogeneity of TBI is also observable at structural brain connectivity of patients.

### 4.3. Revisiting structural plasticity in TBI

Diffuse axonal injury is one of the major characteristics of TBI, which causes disruptions in the connectivity between brain regions [1], leading to cognitive deficits especially in moderate-to-severe cases [56]. Rehabilitation is known to improve cognitive functions of patients [42]. Neuroplasticity, that is, the adaptive changes of structural and functional neural circuitry in terms of molecular, synaptic, and cellular changes, is commonly cited as a potential explanation for the cognitive and functional recovery [57,62]. Although axonal sprouting and functional rewiring post TBI is reported [7,40], the underlying mechanism of change in white matter structural connectivity over time at the network level is still unclear [76].

The significant decline in network similarity of patients relative to healthy controls over time (Fig. 4), may be indicative that the connectivity alterations happening in the network are mainly degeneration in connectivity rather than a recovery. This is in line with consistent neurodegeneration and neuronal loss that is widely reported in the TBI literature, which starts with injury and continues decades post-injury [16,20,32]. An alternative explanation for connectivity alterations in favor of structural recovery could be that the network topology of patients reorganizes to converge to a new normal unlike that of healthy controls to regain the network integrity. The decline of longitudinal change in the similarity of patients among themselves being steeper (Sections 3.3 and SI.1) than that of their similarity relative to healthy controls (Fig. 4), however, contradicts this alternative, further supporting the point that the alterations in the white matter network are not a recovery but a degeneration.

**Figure 4.**
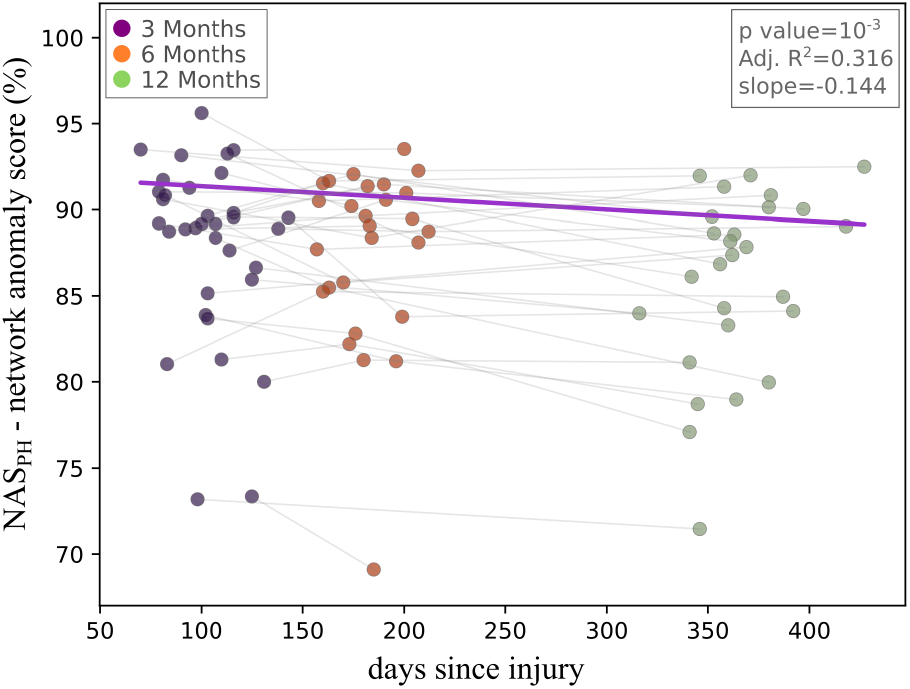
Analysis of change in network anomaly score of patients. Using an LMEM, we evaluated the change in NAS_PH_ score as a function of days since injury, PTA, age, and sex, observing a significant decline in NAS_PH_ with time. This result indicates that the structural network topology of the patients becomes unlike that of healthy controls over time.

In contrast to the decline in their NAS, the cognitive recovery of patients over time (Fig. SI.2) highlights an interesting disparity. When considered together with the lack of a significant association between the rate of change in NAS and cognition (Fig. 5), it can be inferred that the structural changes in the network topology do not directly translate into cognitive recovery. Considering that TBI is a complex disease with multiple, potentially opposing, mechanisms at work simultaneously [67], there might be several reasons for this apparently paradoxical disparity between structural connectivity degeneration and cognitive recovery [16]. One possible explanation is that neuroplasticity happens at the gray matter in terms of axonal sprouting more than white matter plasticity such as myelination. Supporting this perspective, axonal rewiring and sprouting in cortical gray matter are reported to happen in mice post TBI [40]. Several studies on functional MRI, which investigate connectivity of gray matter regions, reported network reorganization after TBI which correlates with cognitive recovery, providing further evidence to that option [7]. Complementing this perspective of synaptic plasticity, another mechanism at play could be that structural connectivity is disrupted at the time of injury, leading to cognitive deficit, due to axonal damage. Although those injured axons do not get repaired and are practically non-functional, some are captured by MRI as healthy fiber tracts connecting brain regions due to the coarse resolution of imaging data. This makes the network topology of a patient look similar to that of a healthy control. As the debris of the damaged axons gets removed from the network, on the other hand, network similarity of patients declines. Since injured axons do not function following the injury, their removal from the network does not have any effect on the cognitive scores of patients as it does not introduce any further disconnectivity into the network.

**Figure 5.**
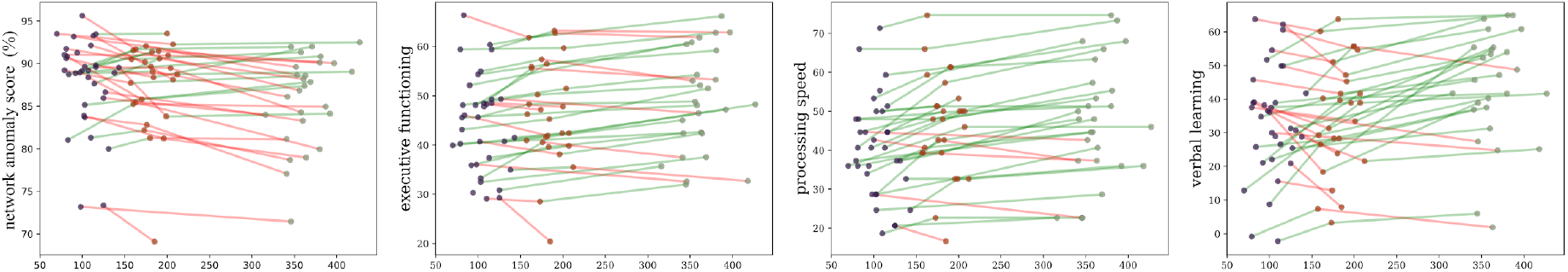
Change in NAS and cognitive scores across time. Plotting individual trajectories of change in NAS_PH_ and cognitive scores for each patient, we observed a steady increase (green lines) for cognitive scores in most cases indicating recovery. In contrast, we observed several cases of decrease (red lines) for NAS_PH_ indicating deviation from normalcy in terms of network topology. Calculating the correlation between the slopes of lines in NAS_PH_ with the slopes of lines in each cognitive score separately, we did not observe any significant relationship. This result indicates that the rate of change in NAS is not associated with cognitive recovery.

We note that the positive correlation between the NAS and EF and PS do not contradict the earlier observations of network similarity declining over time while cognitive scores improve. Since higher NAS values indicate a lesser injury, better cognitive performance would be expected from such individuals as disconnectivity between regions will be lesser. Thus, the negative correlation between injury severity as quantified by PTA and both the NAS and cognitive scores support a positive correlation between the NAS and cognition.

## 5. Limitations, Future Directions, and Conclusions

Although this study investigates a unique longitudinal TBI dataset with dMRI data and cognitive assessment acquired at three timepoints and uses an advanced graph theoretical technique, certain limitations should be acknowledged. First, diffusion MRI is known to have inaccuracies in determining connectivity between regions, such as its limitations in characterizing white matter in complex regions where fibers intersect [33]. In the case of TBI, axonal injury causing the degeneration of one of the crossing fibers, for example, can result in increased FA over the other fiber, which in turn results in increased connectivity between two regions [22]. Since such shortcomings are inherent to dMRI based analysis, the results presented here should be considered accordingly. Second, as typical of TBI studies, statistical significance of our results is limited by the sample size of TBI cohort [69]. Also, our study lacks mild TBI patients, and it should be noted that the results may not translate to a lower injury severity. In order to evaluate the trajectory of structural change in the acute as well as chronic phase of the disease across the injury spectrum, re-evaluation of results presented here on a larger dataset (such as TRACK-TBI, https://tracktbi.ucsf.edu/, [75]) is left as a future work.

In conclusion, our results demonstrate that the structural brain networks of patients with moderate-to-severe TBI differ from those of healthy controls by 3 months and become increasingly different up to 1 year post-injury. It also demonstrates the efficacy of our network anomaly score (NAS) as a principled measure for evaluating severity of diffuse injury, which can have potential uses in creating diagnostic and prognostic biomarkers of the disease when evaluated on larger datasets. Moving forward, we will expand our method to investigate changes in network topology of functional connectivity in TBI patients, in order to explore mechanisms of cognitive recovery with an overall network analysis perspective.

## Acknowledgements

Work of YO, DP, JAA, RV was supported by the National Institutes of Health (RO1 NS096606; YO, DP, JAA, RV). RRD’s work was supported by the Pennsylvania Department of Health, NINDS U01 NS114140, DoD W81XWH-19-2-0002, and W81XWH1910861. The data acquisition for this research was supported by the NIH grant R01-NS065980 (PI: JJK). JJG reports receiving salary support from the National Institute Of Neurological Disorders and Stroke (T32NS091006) and the American Epilepsy Society/Citizens United for Research in Epilepsy (Research and Training Fellowship for Clinicians).

