## Supplementary Materials for "Connectomic Assessment of Injury Burden and Longitudinal Structural Network Alterations in Moderate-to-severe Traumatic Brain Injury"

### SI.1. Mixed effects model analysis of the relationship between network similarity measure, days since injury, injury severity, age, and sex

In order to investigate whether the network organization of patients demonstrates a linear change over time when considered altogether, we evaluated the longitudinal change in their network anomaly score ($NAS_{PH}$ and $NAS_P$ separately). Since the imaging data was not available for each subject across all time points, we used linear mixed effects (LME) analysis with the following model:

$$NAS \sim DSI + PTA + age + sex + (1|subjectID) \qquad \text{(SI.1)}$$

where we estimated NAS as a linear function of the fixed variables days since injury (DSI), post traumatic amnesia (PTA), age of the subject at the first scan, and sex, along with the random variable of subject IDs.

We first considered the relationship for the network similarity of patients relative to healthy controls, and then considered the relationship for the network similarity of patients relative to other patients within the same time point. Results of both linear mixed effect models are reported in Tables SI.1.a and SI.1.b, respectively. Figure SI. 1 shows the decline of $NAS_P$ with time.

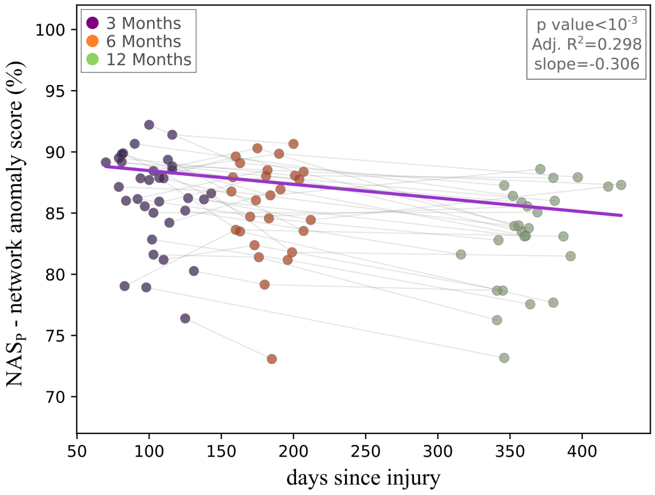

Figure SI.1. Analysis of change in network anomaly score of patients. Using an LMEM, we evaluated the change in $NAS_P$ as a function of days since injury, PTA, age, and sex, observing a significant decline in $NAS_P$ with time. With a steeper slope for the fitted line relative to Fig. 4, this result indicates that the structural network topology of the patients becomes more unlike each other over time compared to their similarity relative to healthy controls. This result also demonstrates that the structural network topology of patients does not converge to a new normal that is different from the healthy controls.

Table SI.1.a. LMEM fit results for evaluating the association between network anomaly score of patients relative to healthy controls ($NAS_{PH}$) and days since injury (DSI), injury severity (PTA), age, and sex (see eqn. SI.1)

| $R^2$ | $p_{DSI}$ | $p_{PTA}$ | $p_{age}$ | $p_{sex}$ | $est_{DSI}$ | $est_{PTA}$ | $est_{age}$ | $est_{sex}$ |
|---|---|---|---|---|---|---|---|---|
| 0.3163 | **0.0009** | **0.0038** | **0.019** | 0.5807 | **-0.144** | **-0.4293** | **-0.3303** | 0.1627 |

Table SI.1.b. LMEM fit results for evaluating the association between network similarity of patients relative to other patients within the same time point ($NAS_P$) and days since injury (DSI), injury severity (PTA), age, and sex (see eqn. SI.1).

| $R^2$ | $p_{DSI}$ | $p_{PTA}$ | $p_{age}$ | $p_{sex}$ | $est_{DSI}$ | $est_{PTA}$ | $est_{age}$ | $est_{sex}$ |
|---|---|---|---|---|---|---|---|---|
| 0.2983 | **$<10^{-4}$** | **0.0073** | 0.066 | 0.6615 | **-0.3061** | **-0.3778** | -0.2444 | 0.1233 |

### SI.2. Analysis of change in cognitive function scores of patients over time

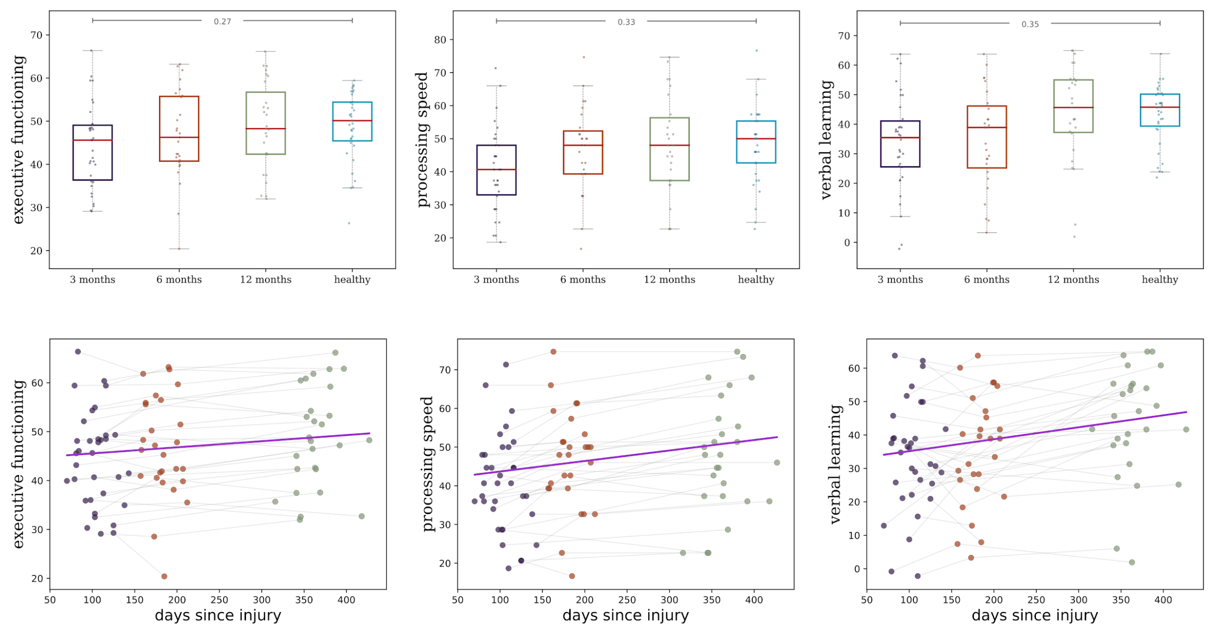

**Figure SI.2.** Analysis of cognitive recovery. (top) We observed patients to have significantly lower cognitive scores compared to healthy controls at 3 months with a medium effect size. At 12 months, however, cognitive scores of patients reach the level of healthy controls. (bottom) Linear mixed effect model reveals a significant improvement in cognitive scores with time 3 to 12 months.

As described in the results section, we observed significant changes in the network organization of patients over time through various analysis methods that we noted above. Potentially, such structural changes in brain network might lead to cognitive and behavioral manifestations. In order to evaluate this, we first investigated whether there is a group difference in cognitive scores between patients and controls. Using Mann-Whitney U test, we observed significant group differences at 3 months for processing speed (ES=0.33, p=0.031) and verbal learning (ES= 0.35, p=0.031), while executive functioning demonstrating difference with a marginal p-value (ES= 0.27, p=0.082) (Fig. SI.2).

We then evaluated whether there exists a significant change in cognitive scores of patients with time using the following LME model:

$$cognitiveScore \sim DSI + PTA + age + sex + (1|subjectID) \quad \text{(SI.2)}$$

where we predicted the three cognitive scores (i.e., EF, PS, and VL) separately as a linear function of DSI, PTA, age, and sex while considering repeated observations of the same subject as the random variable. Results of fitted models are presented in Table SI. 2.

Table SI.2. Linear mixed effects model fit results for evaluating the relationship between cognitive scores and days since injury (DSI), injury severity (PTA), age, and sex (model equation is shown in eqn. SI.2, p-values are FDR corrected for each variable across three models).

| Score | $R^2$ | $p_{DSI}$ | $p_{PTA}$ | $p_{age}$ | $p_{sex}$ | $est_{DSI}$ | $est_{PTA}$ | $est_{age}$ | $est_{sex}$ |
|---|---|---|---|---|---|---|---|---|---|
| Executive Function | 0.433 | $\mathbf{<10^{-5}}$ | $\mathbf{<10^{-4}}$ | 0.548 | 0.391 | 0.013 | -0.295 | -0.077 | 2.325 |
| Processing Speed | 0.464 | $\mathbf{<10^{-6}}$ | $\mathbf{<10^{-4}}$ | 0.279 | 0.344 | 0.027 | -0.379 | -0.189 | 5.624 |
| Verbal Learning | 0.170 | $\mathbf{<10^{-5}}$ | **0.040** | 0.948 | 0.344 | 0.036 | -0.236 | 0.010 | 6.119 |

### SI.3. Mixed effect model analysis of the relationship between cognitive scores and network similarity measure, days since injury, age, and sex

We used a linear mixed effects model to evaluate the relationship between cognitive scores and network similarity score of patients relative to healthy controls, days since injury, age, and sex using the following formulation:

$$cognitiveScore \sim NAS + DSI + age + sex + (1|subjectID) \quad \text{(SI.3)}$$

Results of the fitted model are shown in Table SI.4. We note that we didn't use PTA as one of the predictor variables since we have already shown in Sections 3.2 and 3.3 that network similarity captures injury burden, thus the two scores capture similar phenomena and are collinear.

Table SI.3. Linear mixed effects model fit results for evaluating the relationship between cognitive scores and network similarity score of patients relative to healthy controls (NS), days since injury (DSI), age, and sex (model equation is shown in eqn. SI.3, p-values are FDR corrected for each variable across three models).

| Score | $R^2$ | $p_{NAS}$ | $p_{DSI}$ | $p_{age}$ | $p_{sex}$ | $est_{NS}$ | $est_{DSI}$ | $est_{age}$ | $est_{sex}$ |
|---|---|---|---|---|---|---|---|---|---|
| Executive Function | 0.206 | **$<10^{-3}$** | **$<10^{-4}$** | 0.219 | 0.704 | **0.315** | **0.171** | -0.206 | 0.114 |
| Processing Speed | 0.226 | **0.006** | **$<10^{-4}$** | 0.126 | 0.542 | **0.256** | **0.203** | -0.304 | 0.286 |
| Verbal Learning | 0.141 | 0.086 | $10^{-4}$ | 0.870 | 0.542 | 0.219 | 0.316 | -0.025 | 0.307 |

### SI.4. Evaluation of standard graph theoretical measures over the TBI cohort

In this section, we provide a thorough explanation of the analysis made on standard graph theoretical measures.

First we provide a brief description of each graph theory measure that was used in the study along with references from the TBI literature that report significant changes in the structural connectivity of patients as captured by each measure.

- ***Node betweenness centrality*** quantifies how many times a node appears in the shortest paths between node pairs across the network, with higher values indicating that the node participates in a large number of shortest paths. Reduced betweenness centrality is reported in hub nodes for TBI patients [ref].
- ***Eigenvector centrality*** is a recursively defined score quantifying centrality of a node, where nodes that are connected to nodes with high eigenvector centrality have a higher score. Reduced eigenvector centrality is observed in hub nodes for TBI patients [ref].
- ***Clustering coefficient*** is defined as the fraction of triangles around a node, quantifying the extent of nodes of a graph being clumped together. Higher [ref] as well as lower [ref] clustering coefficient is reported in TBI patients.
- ***Small-worldness*** is defined as the ratio between clustering coefficient and characteristic path length of a network. A network with a score larger than 1 means that it has a small world structure, indicating that, although most nodes are connected to many others directly, each node is connected to the rest of the network through a small number of indirect connections. Higher small-worldness is reported for TBI patients [ref].
- ***Characteristic path length*** is the average shortest path length in a network, where larger values indicate a network with less number of direct connections. Characteristic path length is reported to be larger in TBI patients [ref].
- ***Global efficiency*** is defined as the average inverse shortest path length in the network. Lowered global efficiency is observed in TBI patients [ref].
- Modularity quantifies the strength of a network being divided into modules. Increased modularity is reported in TBI patients [ref].

Our investigation of graph theory measures consist of the following analyses:

a) Relationship between network similarity measure and graph theory measures
b) Relationship between graph theory measures and injury severity, days since injury, age, and sex
c) Relationship between cognitive measures and graph theory measures

In our evaluation, we considered network similarity scores of patients relative to healthy controls. Next, we provide the details of each of the analyses that are listed above.

#### a. Analysis of relationship between network similarity measure and graph theory measures

In our analysis, we first calculated Pearson's correlation between network similarity score and each graph theory score cross sectionally. As shown in Table SI.4.a, no significant correlation was observed.

Table SI.4.a. Pearson's correlation between network similarity score and each graph theory score cross sectionally. Correlation coefficients are reported in each cell with p-values before multiple comparison correction being provided within parenthesis (p-value for each test is equal to 0.928 after FDR correction).

| Measure | **3 Months** | **6 Months** | **12 Months** |
|---|---|---|---|
| Node Betweenness Centrality | -0.2031 (0.2493) | -0.009 (0.9644) | 0.138 (0.4924) |
| Eigenvector Centrality | 0.2529 (0.149) | -0.0934 (0.6429) | 0.1031 (0.6087) |
| Clustering Coefficient | -0.0304 (0.8647) | -0.2091 (0.2953) | -0.0517 (0.7979) |
| Small Worldness | 0.1448 (0.4139) | -0.1524 (0.448) | 0.0867 (0.6674) |

| | | | |
|---|---|---|---|
| Characteristic Path Length | -0.026 (0.884) | 0.3082 (0.1178) | 0.1225 (0.5429) |
| Global Efficiency | -0.0559 (0.7533) | -0.1286 (0.5226) | -0.1447 (0.4715) |
| Modularity | 0.03 (0.8664) | 0.1644 (0.4125) | 0.1069 (0.5956) |

We then used a LMEM to evaluate the relationship between network similarity and graph theory measures using the following model:

$$NAS \sim graphTheoryMeasure + DSI + age + sex + (1|subjectID) \quad \text{(SI.4.a)}$$

which is evaluated for each graph theory measure individually. As shown in Table SI.4.b, no significant relationship was observed.

Table SI.4.b. Linear mixed effects model fit results for evaluating the relationship between network similarity and graph theory measures (GTM), days since injury (DSI), age, and sex (model equation is shown in eqn. SI.4.a, p-values FDR corrected).

| Measure | $R^2$ | $p_{GTM}$ | $p_{DSI}$ | $p_{age}$ | $p_{sex}$ | $est_{GTM}$ | $est_{DSI}$ | $est_{age}$ | $est_{sex}$ |
|---|---|---|---|---|---|---|---|---|---|
| Node Betweenness Centrality | 0.022 | 0.984 | 0.388 | 0.452 | 0.821 | 0.037 | -0.074 | -0.131 | -0.068 |
| Eigenvector Centrality | 0.020 | 0.984 | 0.388 | 0.452 | 0.821 | 0.010 | -0.068 | -0.113 | -0.069 |
| Clustering Coefficient | 0.020 | 0.984 | 0.388 | 0.452 | 0.821 | -0.003 | -0.069 | -0.114 | -0.068 |
| Small Worldness | 0.031 | 0.767 | 0.388 | 0.452 | 0.821 | 0.100 | -0.078 | -0.141 | -0.072 |
| Characteristic Path Length | 0.032 | 0.767 | 0.388 | 0.452 | 0.821 | 0.114 | -0.079 | -0.140 | -0.119 |
| Global Efficiency | 0.032 | 0.767 | 0.388 | 0.452 | 0.821 | -0.122 | -0.063 | -0.159 | -0.117 |
| Modularity | 0.030 | 0.767 | 0.388 | 0.452 | 0.821 | 0.109 | -0.073 | -0.163 | -0.104 |

#### b. Analysis of relationship between graph theory measures and injury severity, days since injury, age, and sex

We did a linear model analysis to evaluate whether graph theory measures can predict injury burden cross sectionally as quantified by PTA. We used the following linear model:

$$PTA \sim GraphTheoryMeasure + age + sex$$

(SI.4.b)

for each graph theory measure individually at each time point. Result of the fitted model is presented in Table SI.4.c. The results indicate a significant relationship only for Node betweenness centrality at 6 months.

Table SI.4.c. Linear model fit results for evaluating whether PTA can be predicted by graph theory measures cross sectionally. (model equation is shown in eqn. SI.4.c, significant relationships are highlighted in bold, p-values are FDR corrected)

| Measure | Adj.$R^2$ | $p_{GTM}$ | $p_{age}$ | $p_{sex}$ | $est_{GTM}$ | $est_{age}$ | $est_{sex}$ |
|---|---|---|---|---|---|---|---|
| **3 Months** | | | | | | | |
| Node Betweenness Centrality | 0.0442 | 0.3197 | **0.042** | 0.9209 | 0.2983 | -0.0173 | 0.4732 |
| Eigenvector Centrality | -0.0287 | 0.8759 | 0.4894 | 0.9765 | 0.0378 | 0.1342 | 0.44 |
| Clustering Coefficient | 0.0174 | 0.4153 | 0.0978 | 0.9765 | 0.225 | 0.0429 | 0.4399 |
| Small Worldness | -0.0091 | 0.5858 | 0.3813 | 0.9841 | -0.1409 | 0.1539 | 0.4294 |
| Characteristic Path Length | 0.0342 | 0.3335 | 0.0822 | 0.9209 | 0.2689 | 0.0229 | 0.339 |
| Global Efficiency | -0.0102 | 0.5858 | **0.042** | 0.9209 | -0.1577 | 0.0526 | 0.3796 |
| Modularity | 0.0091 | 0.4584 | **0.0436** | 0.9209 | 0.2189 | 0.0268 | 0.3558 |

| | | | | | | | |
|---|---|---|---|---|---|---|---|
| **6 Months** | | | | | | | |
| **Node Betweenness Centrality** | 0.5038 | **0.0021** | 0.962 | 0.9841 | 0.6919 | 0.0756 | 0.2702 |
| Eigenvector Centrality | 0.196 | 0.127 | 0.962 | 0.9765 | -0.4255 | 0.138 | 0.4864 |
| Clustering Coefficient | -0.0045 | 0.872 | 0.1848 | 0.9841 | 0.0573 | 0.143 | 0.5039 |
| Small Worldness | 0.0086 | 0.6737 | 0.2224 | 0.9841 | -0.1256 | 0.2018 | 0.5017 |
| Characteristic Path Length | 0.0028 | 0.7328 | 0.4998 | 0.9209 | 0.1022 | 0.1451 | 0.4574 |
| Global Efficiency | 0.0253 | 0.5805 | 0.3813 | 0.9209 | -0.1918 | 0.1187 | 0.3763 |
| Modularity | 0.0715 | 0.3335 | 0.3813 | 0.9209 | 0.2856 | 0.0875 | 0.3872 |
| **12 Months** | | | | | | | |
| Node Betweenness Centrality | 0.1753 | 0.1166 | 0.0822 | 0.9765 | 0.5315 | -0.212 | 0.3191 |
| Eigenvector Centrality | -0.0979 | 0.9471 | **0.0458** | 0.9209 | 0.0163 | -0.0029 | 0.3357 |
| Clustering Coefficient | 0.02 | 0.2877 | 0.0648 | 0.9765 | 0.3608 | -0.1711 | 0.3637 |
| Small Worldness | 0.0234 | 0.2877 | 0.2963 | 0.9841 | 0.3392 | -0.0981 | 0.3363 |
| Characteristic Path Length | 0.0985 | 0.1474 | 0.5688 | 0.9209 | 0.4349 | -0.0611 | 0.1144 |
| Global Efficiency | 0.0308 | 0.2877 | 0.1081 | 0.9209 | -0.3719 | -0.1455 | 0.1848 |
| Modularity | 0.1298 | 0.127 | **0.042** | 0.9209 | 0.534 | -0.2776 | 0.1209 |

#### c. Analysis of relationship between cognitive measures and graph theory measures

We utilized a LMEM analysis to evaluate the relationship between graph theory measures and cognitive scores, using the following formulation:

$$cognitiveScore \sim graphTheoryMeasure + DSI + age + sex + (1|subjectID) \quad \text{(SI.4.c)}$$

where cognitive score is replaced by EF, PS, and VL whole graphTheoryMeasure is replaced by one of the seven measures mentioned early in this chapter, respectively.

Results of the model fit are presented in Table SI.4.d. After FDR correction for multiple comparisons, no significant result was observed for graph theory measures predicting cognitive scores.

Table SI.4.d. Linear mixed effects model fit results for evaluating the relationship between cognitive scores and graph theory measures (GTM), days since injury (DSI), age, and sex (model equation is shown in eqn. SI.4.d, significant relationships are highlighted in bold, p-values are FDR corrected).

| Measure | $R^2$ | $p_{GTM}$ | $p_{DSI}$ | $p_{age}$ | $p_{sex}$ | $est_{GTM}$ | $est_{DSI}$ | $est_{age}$ | $est_{sex}$ |
|---|---|---|---|---|---|---|---|---|---|
| **Executive Function** | | | | | | | | | |
| Node Betweenness Centrality | 0.06 | 0.8336 | **$<10^{-4}$** | 0.45 | 0.9166 | 0.0324 | 0.1345 | -0.2048 | -0.0522 |
| Eigenvector Centrality | 0.0654 | 0.3008 | **$<10^{-4}$** | 0.45 | 0.9166 | 0.0735 | 0.1426 | -0.1741 | -0.0582 |
| Clustering Coefficient | 0.0721 | 0.3008 | **$<10^{-4}$** | 0.6078 | 0.9166 | -0.1335 | 0.1489 | -0.1335 | -0.0544 |
| Small Worldness | 0.062 | 0.4717 | **$<10^{-4}$** | 0.45 | 0.9166 | -0.0605 | 0.1445 | -0.1742 | -0.0482 |
| Characteristic Path Length | 0.0596 | 0.8573 | **$<10^{-4}$** | 0.45 | 0.9166 | -0.0131 | 0.1396 | -0.1878 | -0.0472 |
| Global Efficiency | 0.0634 | 0.8073 | **$<10^{-4}$** | 0.45 | 0.9166 | 0.0477 | 0.136 | -0.1736 | -0.0353 |
| Modularity | 0.0667 | 0.4717 | **$<10^{-4}$** | 0.4058 | 0.9166 | 0.1099 | 0.1354 | -0.241 | -0.0852 |

| | | | | | | | | | |
|---|---|---|---|---|---|---|---|---|---|
| **Processing Speed** | | | | | | | | | |
| Node Betweenness Centrality | 0.1313 | 0.912 | **$<10^{-4}$** | 0.357 | 0.9166 | -0.0079 | 0.2258 | -0.2744 | 0.1445 |
| Eigenvector Centrality | 0.1433 | 0.3008 | **$<10^{-4}$** | 0.357 | 0.9166 | 0.0905 | 0.23 | -0.2572 | 0.138 |
| Clustering Coefficient | 0.1334 | 0.8073 | **$<10^{-4}$** | 0.357 | 0.9166 | -0.0484 | 0.2286 | -0.2571 | 0.1441 |
| Small Worldness | 0.1309 | 0.8573 | **$<10^{-4}$** | 0.357 | 0.9166 | -0.0118 | 0.226 | -0.2746 | 0.1456 |
| Characteristic Path Length | 0.1306 | 0.8336 | **$<10^{-4}$** | 0.357 | 0.9166 | 0.0226 | 0.2231 | -0.283 | 0.1351 |
| Global Efficiency | 0.1502 | 0.4717 | **$<10^{-4}$** | 0.3738 | 0.9166 | 0.1207 | 0.2181 | -0.2342 | 0.1889 |
| Modularity | 0.1321 | 0.8336 | **$<10^{-4}$** | 0.357 | 0.9166 | -0.0342 | 0.2258 | -0.2622 | 0.1548 |
| **Verbal Learning** | | | | | | | | | |
| Node Betweenness Centrality | 0.1108 | 0.3008 | **$<10^{-4}$** | 0.9357 | 0.9166 | -0.1998 | 0.2646 | 0.0551 | 0.2264 |
| Eigenvector Centrality | 0.0827 | 0.4986 | **$<10^{-4}$** | 0.9658 | 0.9166 | 0.0823 | 0.2444 | -0.0115 | 0.2231 |
| Clustering Coefficient | 0.0843 | 0.5187 | **$<10^{-4}$** | 0.9658 | 0.9166 | -0.1185 | 0.2489 | 0.0201 | 0.2281 |
| Small Worldness | 0.0768 | 0.7709 | **$<10^{-4}$** | 0.9357 | 0.9166 | 0.0553 | 0.234 | -0.0453 | 0.2258 |
| Characteristic Path Length | 0.0791 | 0.618 | **$<10^{-4}$** | 0.9357 | 0.9166 | 0.077 | 0.2326 | -0.0481 | 0.1961 |
| Global Efficiency | 0.1014 | 0.4717 | **$<10^{-4}$** | 0.9627 | 0.9166 | 0.1722 | 0.2302 | 0.0324 | 0.2938 |
| Modularity | 0.0755 | 0.8336 | **$<10^{-4}$** | 0.9658 | 0.9166 | -0.0523 | 0.2407 | -0.0067 | 0.245 |
